# Intermittent Hypoxia Drives Early Metabolic Dysfunction in Brown Adipose Tissue

**DOI:** 10.1101/2025.09.18.676807

**Authors:** Nuria Pescador, María Fernández-Cañizares, Yosra Berrouayel, Laura Balaguer, Anna Ulldemolins, Alicia Jurado, Adele Chloe Cardeñosa Pérez, Sandra Pérez, Laura Sánchez, Elena Díaz-García, Alfonso Luis Calle Pascual, Ángela M. Valverde, Isaac Almendros, Germán Peces-Barba, Carolina Cubillos-Zapata, Francisco García-Río, Luis del Peso

## Abstract

**Background:** Obstructive sleep apnea (OSA) is associated with metabolic dysfunction, yet the early impact of intermittent hypoxia (IH)—a defining feature of OSA—on brown adipose tissue (BAT) remains unclear.

**Methods:** We investigated the direct and early effects of IH on BAT using both in vitro and in vivo approaches. Differentiated mouse brown adipocytes were exposed to IH for 48 hours and analyzed for β3-adrenergic signaling, lipolysis, and thermogenic activation. Complementary in vivo studies assessed transcriptomic, morphological, and functional changes in BAT, white adipose tissue (WAT), and liver after 1 or 4 weeks of IH exposure in mice.

**Results:** IH blunted β3-adrenergic signaling in cultured brown adipocytes, leading to reduced phosphorylation of key signaling proteins, impaired lipolytic response, and decreased expression of UCP1. In vivo, BAT exhibited early and sustained transcriptomic rewiring characterized by downregulation of pathways related to fatty acid metabolism, oxidative phosphorylation, and peroxisomal function. These molecular changes were accompanied by abnormal lipid droplet enlargement and a reduced lipolytic response to adrenergic stimulation. In contrast, WAT showed transient gene expression changes, and the liver displayed a delayed and robust transcriptomic response evident only after four weeks of IH exposure. Despite these metabolic alterations, thermogenic responses to cold challenge remained intact in IH-treated mice.

**Conclusions:** BAT is uniquely and rapidly affected by intermittent hypoxia, undergoing functional, molecular, and morphological changes within one week of exposure. These alterations precede systemic inflammation and metabolic dysfunction, positioning BAT dysfunction as an early event in the pathogenesis of OSA-associated metabolic disease.

## INTRODUCTION

Obstructive sleep apnea (OSA) is a common and underdiagnosed sleep disorder characterized by recurrent episodes of partial or complete obstruction of the upper airway during sleep, leading to intermittent hypoxia (IH), fragmented sleep architecture, and increased sympathetic activation (1). Although traditionally viewed as a respiratory condition, OSA is now recognized as a major contributor to metabolic dysfunction, including insulin resistance, dyslipidemia, and obesity (2). The cyclical hypoxia–reoxygenation pattern intrinsic to OSA represents a unique physiological stressor that affects multiple organs, particularly metabolically active tissues.

White adipose tissue (WAT) is especially sensitive to IH. Several studies have shown that IH triggers inflammation, impairs lipid metabolism, and promotes tissue remodeling in WAT (3–7). These changes contribute to the increased cardiometabolic risk observed in OSA patients, independently of obesity, and position WAT as a central player in the metabolic complications associated with sleep-disordered breathing (8).

Alongside WAT, brown adipose tissue (BAT) plays a critical role in energy homeostasis through non-shivering thermogenesis, a process mediated by uncoupling protein 1 (UCP1). This thermogenic function enables BAT to convert stored energy into heat, thereby contributing to cold adaptation and promoting systemic metabolic health, including enhanced insulin sensitivity and energy expenditure (9). The discovery of functional BAT in adult humans (10,11), along with recent evidence linking active BAT to reduced cardiovascular and metabolic disease risk (12) has stimulated interest in its potential as a therapeutic target for obesity and type 2 diabetes (13).

Despite growing interest in BAT, the effects of IH on brown adipocyte function remain poorly understood. A few recent studies have begun to explore this question. For example, Wang et al. (14) reported that prolonged IH exposure (8 weeks) in ApoE□/□ mice activated BAT and promoted atherosclerosis, potentially through increased lipolysis and altered lipid metabolism. Similarly, Dahan et al. (15) found that chronic IH (10 weeks) led to morphological signs of BAT browning, such as smaller lipid droplets and increased vascularization, accompanied paradoxically by mitochondrial dysfunction, inflammation, and insulin resistance. These findings suggest a maladaptive activation of BAT under prolonged IH exposure, which may contribute to the pathophysiology of OSA-related metabolic disease. However, these studies are limited by their long-term exposure models, which make it difficult to discern whether BAT dysfunction is a direct consequence of IH or secondary to systemic metabolic disturbances developing over time. In addition, the early and tissue-specific responses to IH remain largely unexplored. Understanding the early impact of IH on BAT function may provide new insights into the pathophysiological mechanisms linking OSA to metabolic disease, and could help identify early biomarkers or therapeutic targets to prevent or mitigate disease progression.

Here, we aimed to determine how IH directly impairs BAT function and to characterize its early effects on metabolically relevant tissues. Using an in vitro model of differentiated brown adipocytes, we demonstrate that IH blunts β3-adrenergic signaling, reducing both lipolytic and thermogenic responses. Complementary in vivo experiments show that BAT is uniquely and rapidly affected by IH, displaying sustained transcriptomic reprogramming, lipid droplet enlargement, and impaired adrenergic lipolysis after just one week of exposure. In contrast, WAT and liver exhibit either transient or delayed responses to IH. These findings identify BAT as an early target of IH and support a direct role for it in brown adipose tissue dysfunction.

## RESULTS

### Intermittent hypoxia reduces Adrb3 signaling in vitro

To investigate the effect of IH on brown adipose tissue (BAT) function, we exposed in vitro differentiated mouse brown adipocytes to IH for 48 hours and assessed their response to the β3-adrenergic agonist CL316243 (figure 1A). Under these conditions, early signaling events, including phosphorylation of p38 and CREB (figure 1B) and hormone-sensitive lipase (HSL) (figure 1C), were significantly reduced in IH-exposed cells compared to controls.

**Figure 1.**
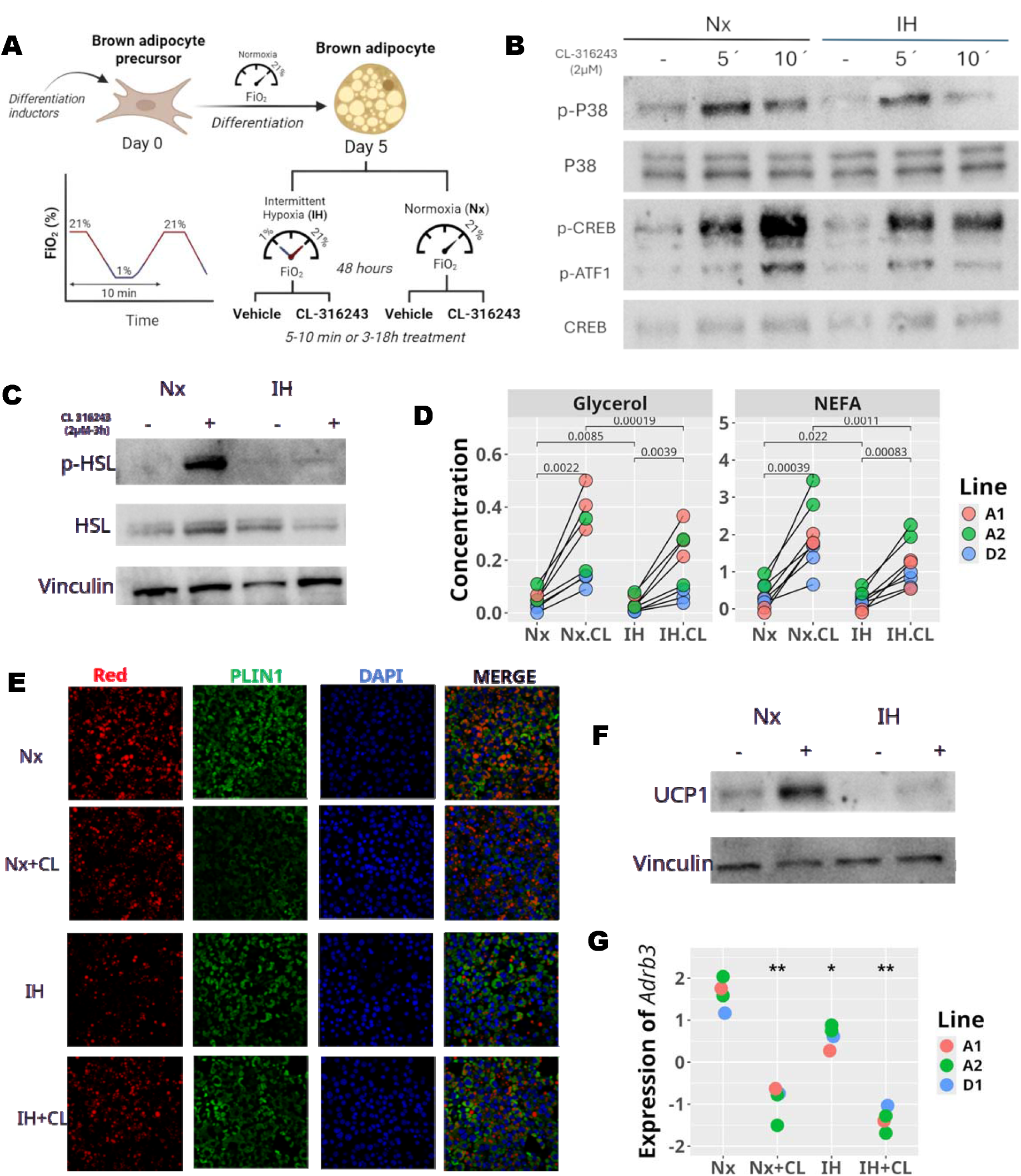
Effect of intermittent hypoxia on cultured brown adipocytes. (A) Schematic of the experimental design. In vitro differentiated brown adipocytes were exposed to IH for 48 hours and then stimulated with the β3-adrenergic agonist CL316243 for either 5-15 minutes (panel B), 3 hours (panel C) or 18 hours (panels D–G). (B–C) Immunoblots showing phosphorylation of the indicated proteins. Images are representative of four independent experiments using three independently generated adipocyte cultures. (D) Glycerol and NEFA levels in supernatants of cells after the indicated treatments were measured by enzymatic assays. Values represent glycerol concentrations (g/L) or NEFA concentrations (mM) in the supernatant. Data are from 8 independent experiments (colored lines), using three different adipocyte cell lines (symbol shapes). Statistical analysis was performed using a repeated-measures ANOVA with cell line as the blocking factor. The analysis revealed a significant effect of treatment (Glycerol, F(3,26) = 17.29, p = 2.2 × 10^−6^; NEFA,F(3,26) = 35.4, p = 2.5 × 10^−9^). Post-hoc pairwise comparisons were conducted using paired t-test, with significance indicated in the figure. (E) Representative micrographs showing Nile Red staining (neutral lipids), Perilipin 1 (PLIN1), and DAPI (nuclei). (F) Immunoblot of UCP1 expression following β3-adrenergic stimulation. Images are representative of four independent experiments. (G) Relative Adrb3 mRNA expression determined by qPCR. Data include four independent experiments (colors) across three adipocyte lines (shapes). Statistical analysis was performed using a repeated-measures ANOVA with cell line as the blocking factor. The analysis revealed a significant effect of treatment (F(3,10) = 58.6, p = 1.2 × 10^−6^). Post-hoc pairwise comparisons with the reference group (Nx) were performed using paired t-tests, with significance indicated in the figure (*p < 0.05; **p < 0.001).

Since CL316243 induces lipolysis in brown adipocytes via phosphorylation-dependent activation of HSL, we next examined whether IH modulates this response. In control cells, CL316243 stimulation led to increased secretion of glycerol and non-esterified fatty acids (NEFAs). However, this lipolytic response was significantly blunted in IH-exposed cells (figure 1D), correlating with the reduced phosphorylation of HSL observed under the same conditions (figure 1C).

To further assess the impact of IH on lipolysis, we performed immunofluorescence for PLIN1 (Perilipin 1) and Nile Red staining. As expected, under normoxic conditions, CL316243-treated cells exhibited smaller lipid droplets and a reduced and redistributed PLIN1 signal, consistent with previous reports (16,17). In contrast, IH-exposed cells treated with CL316243 retained large lipid droplets, and the PLIN1 staining pattern remained unchanged, resembling that of unstimulated cells (figure 1E).

We also evaluated UCP1 expression 18 hours after β3-adrenergic stimulation as an in vitro proxy for thermogenic activation. In line with the impaired lipolytic response, IH-treated cells displayed reduced basal and CL316243-induced levels of UCP1 protein (figure 1F) and mRNA (data not shown).

These findings indicate a defect in β3-adrenergic signaling. Supporting this, we observed that IH exposure significantly decreased basal expression levels of the beta-3 adrenergic receptor, *Adrb3*, mRNA (figure 1H). However, CL316243-induced homologous desensitization of *Adrb3* expression (18) was not affected by IH.

Altogether, these results suggest that IH impairs both the lipolytic and thermogenic responses of brown adipocytes to β3-adrenergic stimulation.

### Intermittent hypoxia induces transcriptomic changes associated with BAT dysfunction

The results described above suggest that IH impairs brown adipocyte function. To further investigate this possibility in a model of OSA, we exposed mice to IH for 1 or 4 weeks and analyzed BAT transcriptome by RNA sequencing (figure 2A). As shown in figure 2B, BAT from IH-exposed animals exhibited several hundred differentially expressed genes (DEGs) at both time points (FDR < 0.05, |log□FC| > 0.5), compared to control animals.

**Figure 2.**
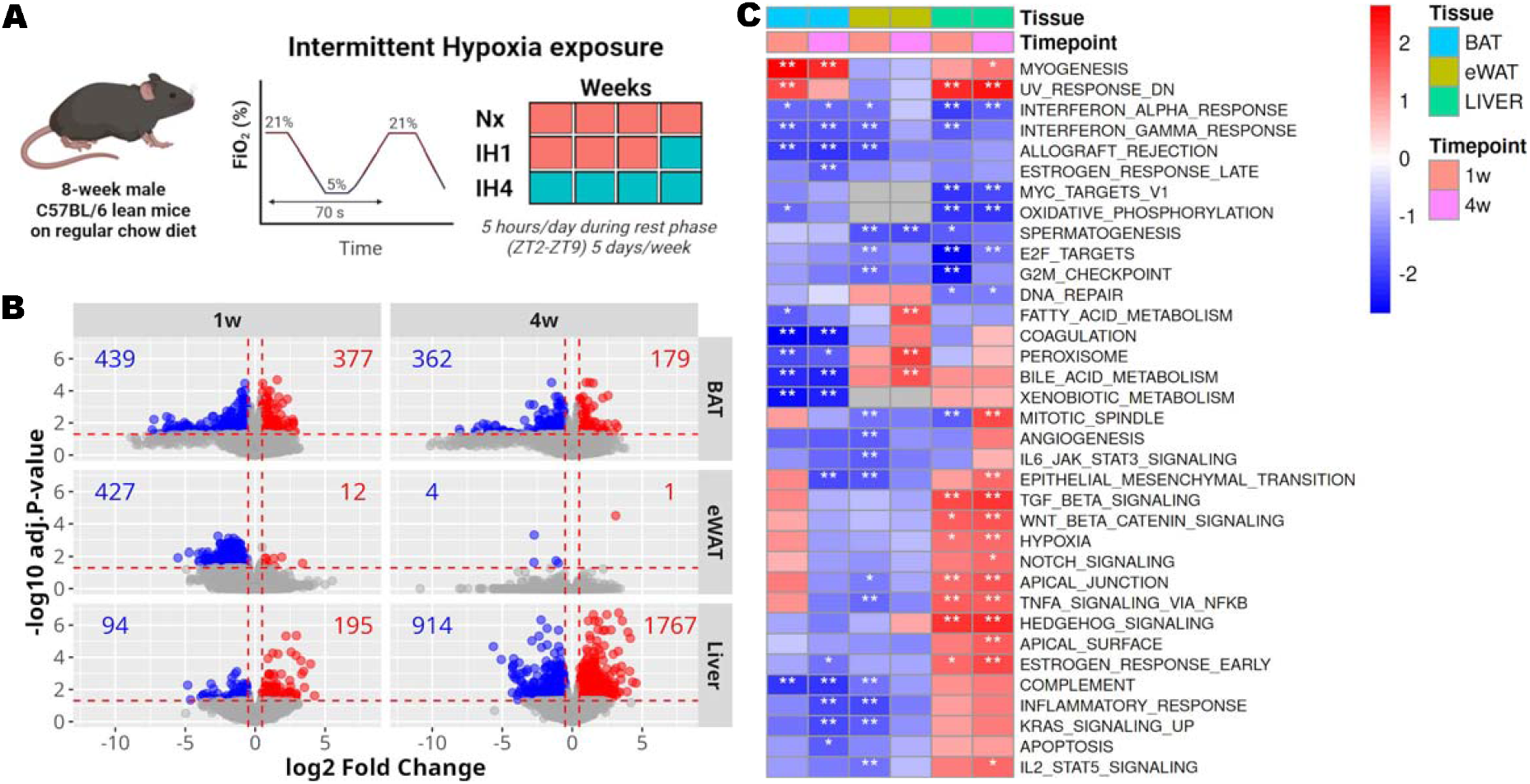
Experimental design and transcriptomic response of metabolically active tissues to intermittent hypoxia. **(A)** Schematic representation of the IH exposure protocol. Adult lean male C57BL/6JRj mice were fed a standard chow diet (55.5% carbohydrates, 16.9% protein, 4.3% fat) and housed under controlled atmospheric oxygen conditions. Mice were exposed in their home cages to either normoxia (Nx) or IH for 5 h/day during their rest phase (ZT2–ZT9). IH consisted of alternating cycles of 21% and 5% inspired oxygen (FiO□) every 70 s (∼50 cycles/h). Nx mice were exposed to normoxia for 4 consecutive weeks, IH 4w mice to IH for 4 consecutive weeks, and IH 1w mice to normoxia for 3 weeks followed by IH for 1 week. Group sizes were 6 for Nx and 7 for IH. **(B)** Volcano plots showing differential gene expression in brown adipose tissue (BAT), epididymal white adipose tissue (eWAT), and liver after 1 and 4 weeks of IH exposure. Each dot represents a gene. The X-axis shows the log□ fold change (IH vs. Nx); the Y-axis indicates statistical significance as –log□ □ (adjusted p-value). Genes showing significant differential expression (FDR < 0.05; |log□FC| > 0.5) are indicated in red for upregulation and blue for downregulation. Group sizes were 6 for Nx and 7 for IH. **(C)** Gene Set Enrichment Analysis (GSEA) using Hallmark gene sets from the MSigDB database. Heatmaps display normalized enrichment scores (NES) for each tissue and time point. Only pathways significantly enriched in at least one condition are shown. *p < 0.05; **p < 0.001; ns = not significant.

We also evaluated the effect of IH on the transcriptome of epididymal white adipose tissue (eWAT) and liver, two organs central to lipid metabolism and implicated in OSA (8,19). While both organs showed transcriptomic responses to IH, their patterns differed from those observed in BAT. In WAT, the response was transient, with a marked reduction in the number of DEGs by 4 weeks (figure 2B). In contrast, the liver displayed a delayed yet robust response: minimal changes were observed at 1 week, but thousands of genes were differentially expressed by 4 weeks of IH exposure (figure 2B).

The temporal dynamics of gene expression changes were further assessed by comparing DEGs at 1 and 4 weeks. In BAT and liver, transcriptomic alterations were highly correlated over time (supplementary figure 1; BAT: slope = 0.97, R^2^ = 0.90; liver: slope = 1.25, R^2^ = 0.91), indicating sustained transcriptional responses. In contrast, WAT showed poor correlation between time points (slope = 0.33, R^2^ = 0.28), consistent with a transient effect of IH. These findings highlight BAT as an early and persistently responsive tissue to IH, in contrast with the delayed hepatic response and the short-lived changes in WAT.

To investigate the functional implications of IH-induced transcriptomic changes in BAT, we performed gene set enrichment analysis (GSEA). As shown in figure 2C, several biological processes were significantly repressed, including pathways critical for BAT function such as fatty acid metabolism, oxydative phosphorylation and peroxisome activity. These findings suggest that IH compromises key metabolic functions of BAT. However, the quantification of mtDNA indicate that the number of mitochondria are not altered in mice exposed to 1 or 4 weeks of hypoxia ruling out as a possible cause of these metabolic defects (supplementary figure 2).

Together, these results indicate that BAT is uniquely sensitive to IH, exhibiting early, sustained transcriptomic reprogramming associated with impaired metabolic and endocrine functions.

### Intermittent hypoxia leads to lipid accumulation and defective lipolysis in BAT

The transcriptomic analysis are suggestive of BAT dysfunction. In agreement, the BAT from mice exposed to IH for 1 or 4 weeks showed cells containing few large lipid droplets (LD), contrasting with the normal morphology, comprising multiple small multilocular lipid deposits, observed in control animals (figure 3A). The morphology was clearly observed after just one week of treatment and still present after four weeks. The analysis of lipid droplet size distribution across different regions of the organ confirmed that animals exposed to IH had a larger proportion of large LD resulting in a right-skewed distribution that was consistent in all the mice included in the study (figure 3B). The integration of all the data across animals indicate that that the distribution of LD in control animals is significantly different to that observed after 1 week (permutation test, z-score = 74.82, p<2.2e-16) or 4 weeks of exposure to IH (permutation test, z-score = 46.73, p<2.2e-16). In agreement, the number of LD of very large size was significantly higher in IH-treated animals that in controls (figure 3C) at both 1 week (chi-squared test, X^2^ = 14115, df = 1, p-value < 2.2e-16; odd-ratio = 3.32) and 4 weeks (chi-squared test, X^2^ = 3746.2, df = 1, p-value < 2.2e-16; odd-ratio = 1.95) of treatment. The effect of IH on lipid accumulation in BAT, was confirmed in several independent experiments (supplementary figure 3). Although the kinetics of accumulation varied across different experiments, with some showing higher accumulation at 1 week while in other the maximum was observed after 4 weeks, in almost all the cases IH resulted in altered BAT morphology showing LD of increased size observable during the whole duration of the experiment. In contrast to BAT, WAT morphology (figure 3A) and the distribution of LD sizes (figure 3B) was similar in IH-treated and control mice. Although, there was a small increase in the number of LD o very large size after one week of treatment (figure 3C), it did not reached statistical significance (chi-squared test, X^2^ = 3.0246, df = 1, p-value = 0.08201; odd-ratio = 1.25) and, by four weeks, the number of very large LD was smaller than in control animals (chi-squared test, X^2^ = 11.236, df = 1, p-value = 0.0008021; odds-ratio = 0.56).

**Figure 3.**
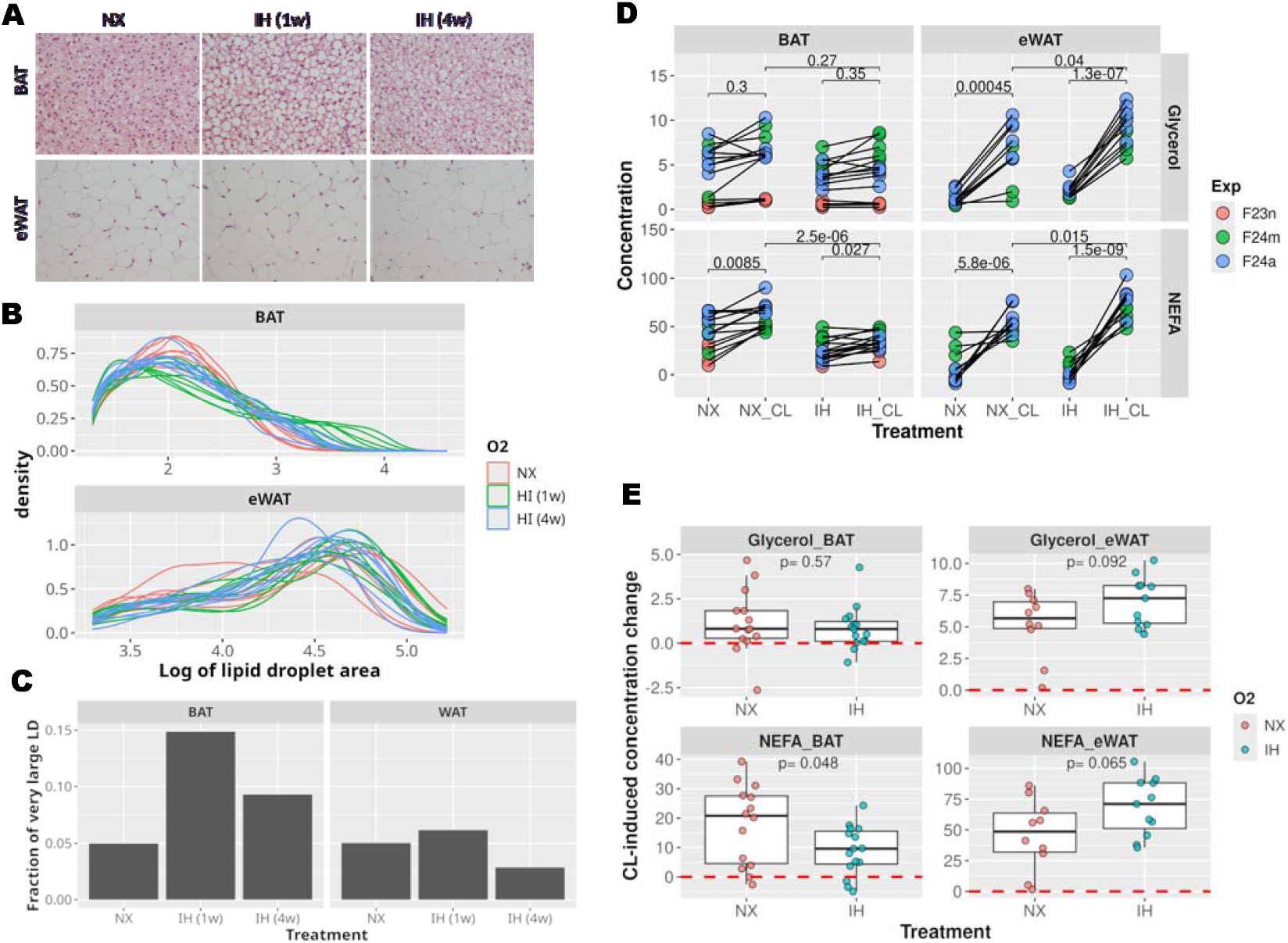
Lipid accumulation and lipolytic response in mice exposed to IH. (A–C) Mice were exposed to IH for 4 weeks, or for 1 week following 3 weeks of normoxia (IH3). Control mice (NX) were maintained under normoxia for 4 weeks. All animals were euthanized on the same day. Group size was 8 for all conditions. (A) Representative hematoxylin-eosin stained paraffin sections of BAT and eWAT. (B) Lipid droplets (LDs) were quantified using the Fiji plugins AdipoQ (BAT) and Adiposoft (eWAT). LD area was measured from images (in pixels^2^) and plotted as log□□(area). Area distributions for individual mice are shown as line plots. (C) Using the LD size distribution from control animals, the 95th percentile cutoff was determined. Bars indicate the fraction of LDs above this threshold in each group. (D–E) Ex vivo lipolysis assay: paired BAT and eWAT fragments from IH- or Nx-treated animals were stimulated with CL316243 or vehicle. (D) Glycerol and NEFA levels in supernatants were measured by enzymatic assays. Values represent glycerol concentrations (g/L) or NEFA concentrations (mM) in the supernatant, normalized to tissue weight (g). Data from paired tissue fragments are connected by lines. Statistical analysis was performed using a repeated-measures ANOVA with exp as the blocking factor. The analysis revealed a significant effect of treatment (BAT/Glycerol, F(3,52) = 5.1, p = 0.003614; BAT/NEFA, F(3,52) = 36.09, p = 7.9 × 10^−10^; eWAT/Glycerol, F(3,37) = 54.4, p = 1.2 × 10^−13^; BAT/NEFA, F(3,37) = 55.5, p = 9.1 × 10^−14^). Post-hoc pairwise comparisons were conducted using paired t-test, with significance indicated in the figure. Total group sizes per condition ranged from 4 to 6 animals per experiment, with 10–15 animals overall across the three independent experiments. (E) Change in metabolite levels following stimulation (CL316243 – vehicle) shown for each tissue. Differences between IH and Nx groups assessed by Student’s t-test and p-values are indicated in the figure. Total group sizes per condition ranged from 4 to 6 animals per experiment, with 10–15 animals overall across the three independent experiments.

IH-treated mice showed a significantly diminished weight gain as compared with control animals (supplementary figure 4), ruling it out as the cause of accumulation of lipids in BAT. The decreased weight gain has been consistently observed across multiple studies and with different IH protocols (15,20–22) and, although we could not assess food intake, other works have demonstrated that the reduced weight gain in not due to reduced food intake (21) and we did not observed differences in leptin expression in BAT nor eWAT (data not shown).

The observed lipid accumulation can be a consequence of increased accumulation or decreased lipolysis. To distinguish between these possibilities we dissected the BAT and eWAT from animals exposed to IH for 1 week and treated them ex-vivo with the β3-adrenergic agonist CL316243 or vehicle. As expected, beta-adrenergic stimulation produced the release of glycerol and NEFA in both eWAT and BAT (figure 3D). However, while the response of eWAT from IH-treated animals was similar to controls, BAT showed a reduced response that reached statistical significance in the case of NEFA (figure 3E). These results are in agreement with in vitro data (figure 1D) and suggest that abnormal lipid accumulation observed in BAT could be, at least in part, due to reduced lipolytic response to adrenergic stimulation.

### Intermittent hypoxia does not alter the thermogenic response of BAT

The results described above suggest a potential dysfunction of BAT. To assess whether IH impairs BAT thermogenic function, we implanted temperature-sensing RFID transponders subcutaneously in the interscapular region and monitored core body temperature throughout the IH exposure protocol. As shown in figure 4A, the IH-treated group exhibited a slightly lower body temperature compared to controls; however, the difference was modest and did not reach statistical significance. The largest difference was observed on the final day of treatment (day 8), with a mean difference of 0.8□°C (SE = 0.36□°C; 95% CI: −0.01 to 1.61).

**Figure 4.**
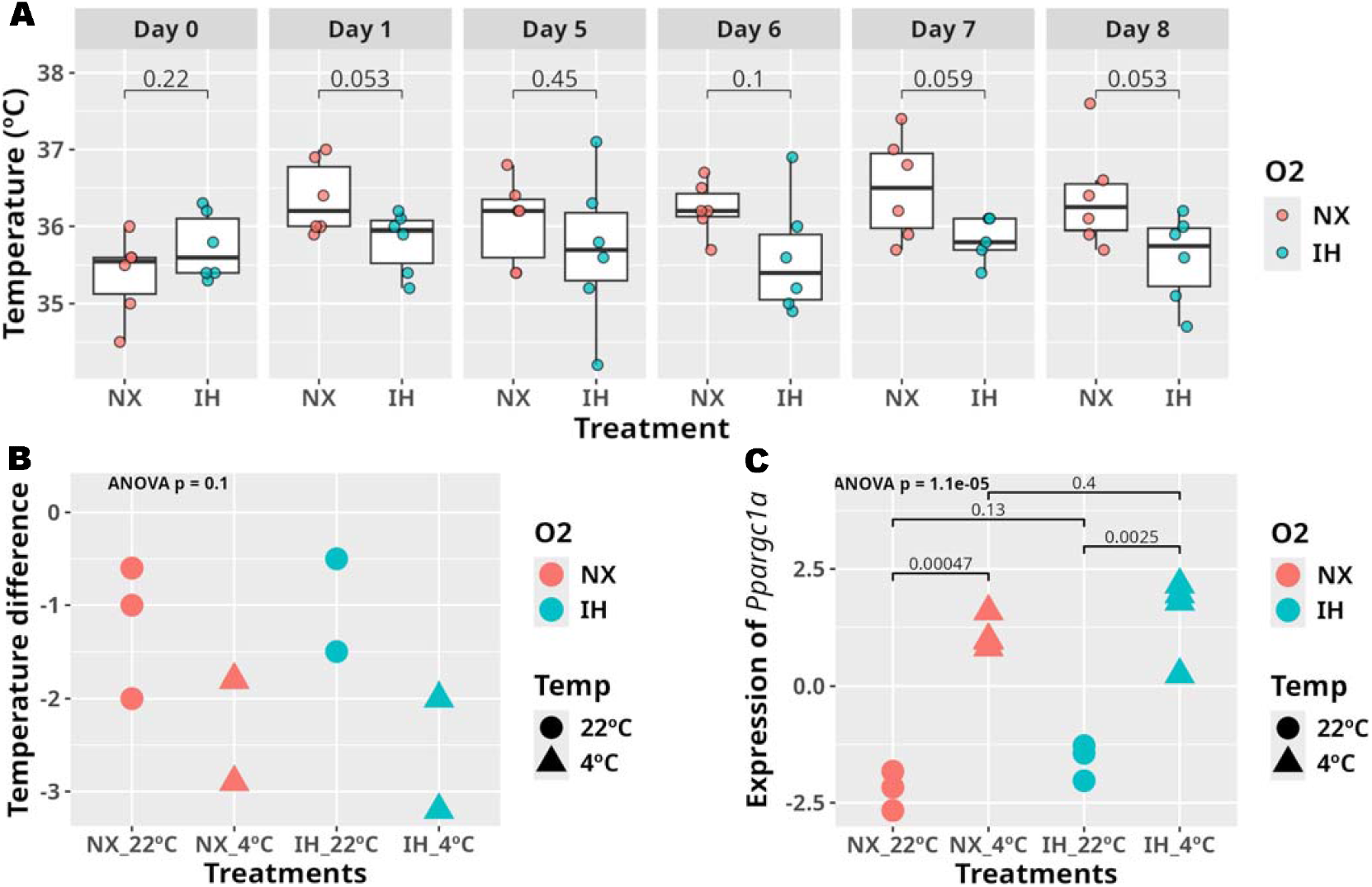
Effect of intermittent hypoxia on core body temperature and thermogenesis. (A) Core body temperature recorded daily at ZT1 during the 1-week IH or control protocol. Statistical significance between groups was assessed using Student’s t-test. Group sizes were 5 for Nx and 6 for IH. (B) Change in body temperature following 16-hour exposure to cold (4□°C) or room temperature (22□°C), measured as post-exposure minus pre-exposure values. No significant difference observed (ANOVA, F(3,6) = 3.224, p = 0.103). (C) Ppargc1a expression in BAT following cold exposure, shown as log□-scaled relative expression normalized to the mean across all samples. Statistical analysis was performed using one-way ANOVA (F(3,24) = 29.64, p = 1.1 × 10^−5^), followed by post-hoc t-tests as indicated.

In addition to basal temperature, we evaluated the cold-induced thermogenic response. After one week of IH or normoxia, animals were exposed to cold (4□°C) or room temperature (22□°C) for 16 hours. Cold exposure resulted in a slight decrease in core body temperature relative to baseline in both groups. However, the magnitude of this reduction was similar between normoxic and IH-exposed animals (figure 4B), and the difference was not statistically significant (ANOVA, p = 0.1).

To further characterize this response, we analyzed the expression of the transcriptional coactivator Ppargc1a, which encodes PGC1α, a key regulator of the molecular program that promotes thermogenesis in BAT in response to cold (23). Consistent with this role, cold exposure induced a robust increase in Ppargc1a expression in both control and IH-treated mice (figure 4C), indicating a comparable molecular response.

Taken together, these results indicate that a one-week exposure to IH does not significantly impair the thermogenic capacity of BAT.

## DISCUSSION

Mounting evidence suggests that the metabolic complications observed in OSA patients are mediated, at least in part, by the effects of IH on key metabolic organs such as the liver and adipose tissues. Within this context, brown adipose tissue (BAT) has received comparatively little attention, despite its central role in systemic energy homeostasis. Moreover, most existing studies have focused on the consequences of prolonged IH exposure, making it difficult to distinguish direct effects of hypoxia from secondary responses to systemic dysfunction. In this study, we addressed this gap by analyzing both the early in vivo responses to IH in BAT and the direct effects of IH on cultured brown adipocytes. Our findings complement and extend previous work, offering novel insights into BAT as an early and direct target of IH.

Our in vitro experiments demonstrate that IH exerts a direct and pronounced effect on brown adipocytes. Specifically, 48 hours of IH exposure markedly attenuated the response to β3-adrenergic stimulation, as evidenced by reduced phosphorylation of key signaling proteins, impaired lipolysis, and decreased expression of UCP1, a core thermogenic effector. In support of a direct action of IH, our in vivo analysis revealed that BAT undergoes clear morphological and functional alterations after just one week of exposure. These changes included the accumulation of large lipid droplets and a blunted lipolytic response to adrenergic stimulation. Gene set enrichment analysis showed that IH suppressed pathways related to fatty acid metabolism, oxidative phosphorylation, and peroxisomal activity, suggesting that diminished lipid catabolism contributes to lipid droplet accumulation.

Transcriptomic profiling revealed that BAT is uniquely and persistently responsive to IH, showing substantial gene expression changes as early as one week into treatment, with these changes sustained through four weeks. In contrast, WAT exhibited a transient response, and the liver showed a delayed but robust transcriptomic shift only after four weeks. These temporal dynamics suggest that IH first targets adipose tissues, particularly BAT, and that hepatic dysfunction may be a downstream consequence of earlier adipose alterations. This interpretation aligns with prior studies reporting that metabolic and inflammatory changes in the liver emerge only after prolonged IH exposure (21,22).

Macrophage-driven inflammation in adipose tissue and liver contributes to insulin resistance and systemic metabolic dysfunction (24,25). While long-term IH has been linked to inflammation in WAT (5), liver (22), and BAT (15), our short-term experiments revealed significant downregulation of inflammatory gene signatures in BAT and WAT, with only a mild, statistically non-significant induction in the liver. Consistent with these transcriptomic data, we found no significant differences in immune cell infiltration between control and IH-treated mice (data not shown). These findings indicate that BAT inflammation is a late event in the IH response. Furthermore, in agreement with studies showing that robust inflammatory responses and insulin resistance typically appear after 8–16 weeks of IH (15,22), our glucose and insulin tolerance tests showed no significant differences between control and IH-treated mice after four weeks (data not shown). Altogether, the morphological and transcriptional changes observed in BAT likely represent early events that precede systemic metabolic dysfunction and inflammation.

Interestingly, Hedgehog signaling was already induced in the liver after just one week of IH exposure. Although typically quiescent in the adult liver, Hedgehog signaling is reactivated in response to injury and correlates with the severity of tissue damage (26). Its early induction may therefore serve as a sensitive marker of IH-induced hepatic stress and a harbinger of later pathology.

With regard to underlying mechanisms, our in vitro data revealed that IH downregulates *Adrb3*, the gene encoding the β3-adrenergic receptor, offering a potential explanation for the impaired adrenergic responsiveness observed. Although we examined the expression of Tribbles pseudokinases, known repressors of *Adrb3* transcription (18), we did not detect changes in *Trib1, Trib2*, or *Trib3* under our experimental conditions (data not shown). Further studies will be required to delineate the molecular pathways by which IH suppresses *Adrb3* expression in brown adipocytes. Notably, in contrast to the in vitro findings, *Adrb3* expression in BAT was not significantly altered in vivo, and thermogenic responses to cold exposure remained largely preserved. These discrepancies suggest the involvement of systemic or compensatory mechanisms in vivo. This divergence highlights a limitation of our study and underscores the need for further investigation to reconcile context-dependent effects of IH.

In conclusion, our findings demonstrate that BAT is an early and primary target of IH, undergoing functional, morphological, and transcriptomic alterations after just one week of exposure. These changes precede the onset of inflammation or systemic metabolic dysfunction and include impaired lipolysis, lipid accumulation, and suppression of key metabolic pathways. Our in vitro data support a direct effect of IH on brown adipocytes, mediated at least in part by repression of β3-adrenergic signaling. Collectively, these results identify BAT dysfunction as a potential initiating event in the cascade of metabolic disturbances associated with OSA and emphasize the importance of early interventions aimed at preserving BAT function in affected individuals.

## MATERIALS AND METHODS

### Generation of brown preadipocyte cell lines and differentiation

Brown preadipocytes were isolated from the interscapular BAT of 20-day old suckling mice and immortalized as previously described (27). In this study, we used several preadipocyte cell lines derived from independent isolation and immortalization procedures. Immortalized preadipocytes were maintained in DMEM supplemented with 100 U/mL penicillin, 100μg/mL streptomycin, 10mM HEPES, and 10% FBS.

For differentiation, immortalized brown preadipocytes were grown in DMEM supplemented with 10% fetal bovine serum (FBS), 20nM insulin and 1nM triiodothyronine (T3) (differentiation medium, DM) until reaching confluence. Next, the cells were cultured for 2 days in induction medium (IM) consisting of DM supplemented with 0.5μM dexamethasone, 0.125μM indomethacin, 1μM rosiglitazone and 0.5mM isobutyl-methyl-xanthine (IBMX). Then, cells were cultured in DM until day 5 in which they exhibited a fully differentiated phenotype.

### Intermittent hypoxia in vitro model

Brown adipocytes at day 5 of differentiation were exposed to IH for 48 hours or left under stardard normoxic culture conditions. For IH exposure plates were placed in an incubation chamber attached to an external oxygen/nitrogen computer-driven controller using BioSpherix OxyCycler C42 (Redfield, NY, USA), a system that generates periodic changes in oxygen concentrations and controls air gas levels in each chamber, while individually maintaining CO2. The IH model cycled oxygen in the medium at 1% for 2 min, followed by 20% for 10 min, with CO2 maintained at 5%. Cells were also cultured under normoxia conditions (21% oxygen, 5% CO2) for the control group.

### Lipolysis assay in brown adipocytes

Brown adipocytes were stimulated for 18h with DMEM-5% FBS and then incubated with Krebs– Ringer Modified Buffer (KRB) (118.5mM of NaCl, 4.75mM of KCl, 1.92mM of CaCl2, 1.19mM of KH4PO4, 1.19mM of MgSO4(H2O)7, 25mM of NaHCO3, 10mM of HEPES, 6mM ofd-glucose, 4% BSA, and pH 7.4) and 2μM CL316243 was added for the last 4h. Supernatants were collected and the amount of glycerol was quantified spectrophotometrically at 540nm with a Free Glycerol Reagent colorimetric kit (12812, BioSystems, Napa, CA, USA). The amount of glycerol was determined using a commercial standard solution. NEFA quantification was performed using the colorometric kit (Ref. 517982 And 517984, Palex Medical, Barcelona, Spain) following the manufacturer’s recommendations.

### Immunofluorescence analysis in brown adipocytes

Brown preadipocytes were differentiated on glass coverslips and then subjected to IH or normoxia for 48h according to the protocol described above. To visualize lipid droplets the cells were fixed in 4%-paraformaldehyde (PFA) and stained with Nile Red solution (0,5 mg/ml in DMSO) (Sigma-Aldrich, Darmstadt, Germany) for 10 min. Then, cells were washed twice with PBS and processed for immunofluorescence. Briefly, anti-PLIN1 D1D8 XP (#9349, Cell Signaling, MA, USA) primary antibody was applied for 18h at 4°C in PBS-1% BSA at 1/100 dilution. The secondary antibody used was Alexa 488 goat anti-mouse (Invitrogen, Waltham, MA, USA) at 1/500 dilution. Cell nuclei were counterstained with DAPI (Sigma-Aldrich, Darmstadt, Germany). Immunofluorescence was examined using an LSM 700 confocal microscope (Zeiss, Oberkochen, Germany).

### Preparation of protein cell extracts and Western blot

Cells were scraped off in ice-cold PBS, pelleted by centrifugation at 4000×g for 10min at 4°C and resuspended in buffer containing 10mM Tris-HCl, 5mM EDTA, 50mM NaCl, 30mM disodium pyrophosphate, 50mM NaF, 100μM Na3VO4, 1% Triton X-100, 1mM PMSF and protease inhibitor cocktail (Complete ULTRA Tablets, Roche) pH 7.6 (lysis buffer). Cell lysates were clarified by centrifugation at 12,000×g for 10min at 4°C twice. Protein content was determined by the Bradford method, using the Bio-Rad reagent and bovine serum albumin (BSA) as the standard. After SDS-PAGE, proteins were transferred to Immobilon membranes (Merk-Millipore), blocked using 3% BSA in 10mM Tris-HCl, 150mM NaCl pH 7.5, and incubated overnight with the antibodies indicated in 0.05% Tween-20, 10mM Tris-HCl, 150mM NaCl pH 7.5. Immunoreactive bands were visualized using the ECL Western blotting protocol (Bio-Rad, Hercules, CA, USA).

### RNA Extraction and real time quantitative PCR

Total RNA extraction and purification were performed using NucleoSpin RNA Plus Kit (Marcherey Nagel, Düren, Nordrhein-Westfalen, Germany) following the manufacturer’s instructions. Complementary DNA (cDNA) was synthesized by reverse transcription from 250ng of RNA using M-MLV reverse transcriptase protocol (M1701, Promega, Madrid, Spain). Real time quantitative PCR was carried out with the Power SYBR Green PCR Master Mix (Applied Biosystems, Waltham, MA, USA). PCR amplifications were performed on the StepOne Realtime PCR System (Applied Biosystems, Waltham, MA, USA). Data were analyzed with StepOne software version v2.1, and expression levels were calculated using ΔΔCt relative to mean expression across conditions. Specifically, for each gene, Ct values were centered by subtracting the mean Ct across all samples, so that lower Ct values (indicative of higher expression) yield positive ΔCt values. Relative expression (ΔΔCt) was calculated by subtracting the centered Ct value of the housekeeping gene from that of each gene of interest for each condition. The resulting values represent log2-scaled relative expression, normalized to the mean expression across samples. Primers used in this study: *Adrb3* (TCCACCGCTCAACAGGTTTG and CTGGATCTTCACGGCCCTTC), *Ppargc1a* (GTCATTCGGGAGCTGGATGG and CAACCAGAGCAGCACACTCT), *Nd1* (CTAGCAGAAACAAACCGGGC and CCGGCTGCGTATTCTACGTT), *Mt16* (CCGCAAGGGAAAGATGAAAGAC and TCGTTTGGTTTCGGGGTTTC), *Hk2* (GCCAGCCTCTCCTGATTTTAGTGT and GGGAACACAAAAGACCTCTTCTGG) and *Rplp0* (GCTTTCTGGAGGGTGTCCG and ACGCGCTTGTACCCATTGAT).

### Animal handling and ethics approval

Eight-week-old male C57BL/6JRj mice (Charles River, Wilmington, MA, USA) were housed under standard conditions with ad libitum access to standard chow and water, on a 12-hour light/dark cycle. Throughout the experiment, mice were maintained on a regular chow diet and randomly assigned to either IH or normoxic control groups.

IH exposure was conducted for 5 hours daily during the animals’ rest phase (from 9:00 AM [ZT2] to 2:00 PM [ZT9]). During these periods, mice were placed in specialized chambers with alternating inflows of 21% O□ and N□, allowing oxygen levels to cycle between 21% and 5% FiO□. Each hypoxia-reoxygenation cycle lasted approximately 70 seconds, achieving ∼50 cycles per hour. Oxygen levels inside the chambers were continuously monitored. Control (Nx) mice were exposed to identical airflow cycles using room air to control for noise and turbulence caused by gas exchange.

Mice were sacrificed by cervical dislocation after either 1 or 4 weeks of exposure. Tissues were rapidly collected and snap-frozen in liquid nitrogen for subsequent analyses.

All animal procedures conformed to the European Directive 2010/63/EU and were approved by the Comunidad Autónoma de Madrid (PROEX 209/19).

### Ex vivo lipolysis assay

Ex vivo lipolysis studies were performed in 8-10 week old male mice. Mice were exposed to conditions of intermittent normoxia or hypoxia for 1 or 4 weeks before being euthanized by cervical dislocation. Ex vivo lipolysis was assessed in white adipose tissue extracted from the epididymal area and brown adipose tissue extracted from the interscapular area following the protocol previously described by Bridge-Comer PE et al (28).

### Histological analysis and immunostaining

Hematoxylin and eosin (H&E) staining was performed in paraffin sections of BAT from mice maintained under IH or normoxia for 1 or 4 weeks according to the protocol indicated above. BAT tissue was fixed in 4% paraformaldehyde (PFA, 16005, Sigma–Aldrich, Darmstadt, Germany) for 24h, washed twice with PBS, dehydrated with ascending ethanol solutions, incubated with xylene, and then embedded in paraffin. Blocks were cut into 5μm sections. Prior to H&E staining, the sections were deparaffinized in xylene and hydrated in descending ethanol solutions and distilled water. The slides were stained with Mayer’s hematoxylin (MHS32-1L, Sigma–Aldrich, Darmstadt, Germany) for 15–20min and eosin (1.15935.0025, Merck, Darmstadt, Germany) for 1min. After the slides were dried and mounted, the images were captured with an Axiophot light microscope (Zeiss, Oberkochen, Germany) using a 40× objective.

### RNA-seq analysis

Immediately after treatments animals were euthanized and their organs were frozen in liquid nitrogen.

Tissues were mechanically disrupted using a Potter-Elvehjem homogenizer and RNA was extracted as indicated above. Library preparation and sequencing were performed by Novogene (Munich, Germany) using the Illumina NovaSeq 6000 platform, generating 150-bp paired-end reads. Raw reads were quality-checked with FASTQC (v0.11.8), then aligned to the mouse reference genome (mm10) using HISAT2 (v2.1.0) with default parameters in paired-end mode. Alignment files (SAM) were converted to BAM and coordinate-sorted using SAMtools (v1.6). Gene-level quantification was performed with HTSeq-count (v0.11.3) using mm10 GTF annotations. Genes with low expression were filtered using the filterByExpr function from the edgeR package (R v4.3.3). Differential expression analysis was conducted with the limma-voom pipeline, and p-values were adjusted for multiple testing using the Benjamini–Hochberg method.

### Statistical analysis

Statistical analyses were carried out using R 4.4.1 (29). Tests applied and their results are shown in figure legends. For the permutation test we assessed differences in the distributions of lipid droplet sizes between the condition and control groups using a permutation test. First, we constructed histograms with 50 bins for each group. For every bin, we calculated the absolute difference in bin height between the two histograms and summed these differences to obtain a test statistic. Next, we performed 1,000 random permutations by shuffling the droplet size values and reassigning them to either the condition or control group. For each permutation, we recalculated the test statistic, thereby generating a null distribution. Finally, we compared the observed test statistic to this null distribution to compute a z-score and corresponding p-value, indicating the significance of the observed difference.

## Supporting information

Text for supplemental Figures

## ACKNOWLEDGMENTS

We thank Dr. Antonio Castrillo for reagents and stimulating discussions about the results presented in this work. We are grateful to Soledad Montalbán and Oscar Sánchez from the CNB Histology Facility at CNB-CSIC (Spain) for histological preparation of biological samples.

## DATA AVAILABILITY

All raw RNA sequencing data generated and analyzed in this study have been deposited in the NCBI Gene Expression Omnibus (GEO) under accession number GSE276218. The data will be made publicly available upon publication of the manuscript.

