## Supplemental Figures for "Intermittent Hypoxia Drives Early Metabolic Dysfunction in Brown Adipose Tissue"

### Slide 1
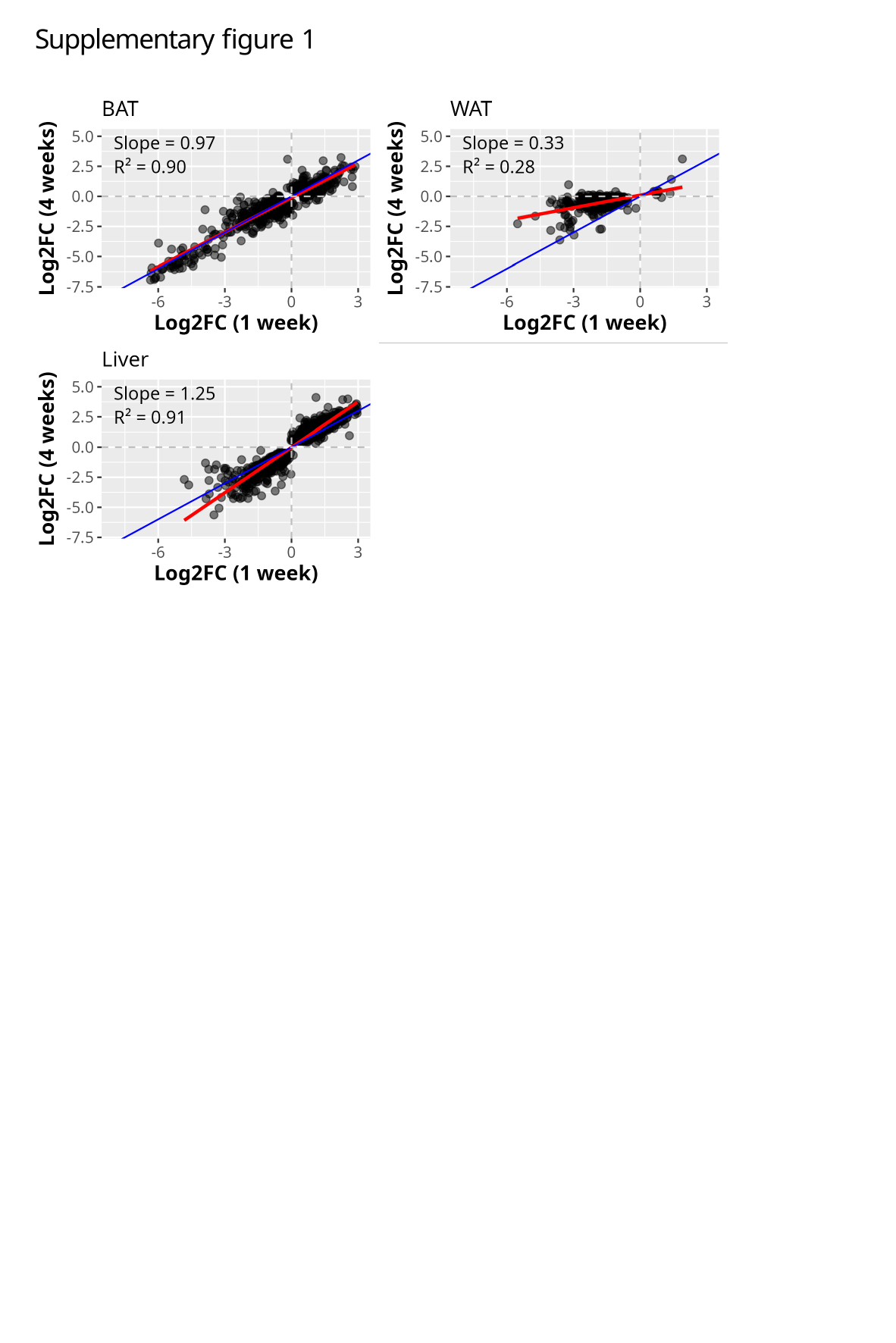

Supplementary figure 1

### Slide 2
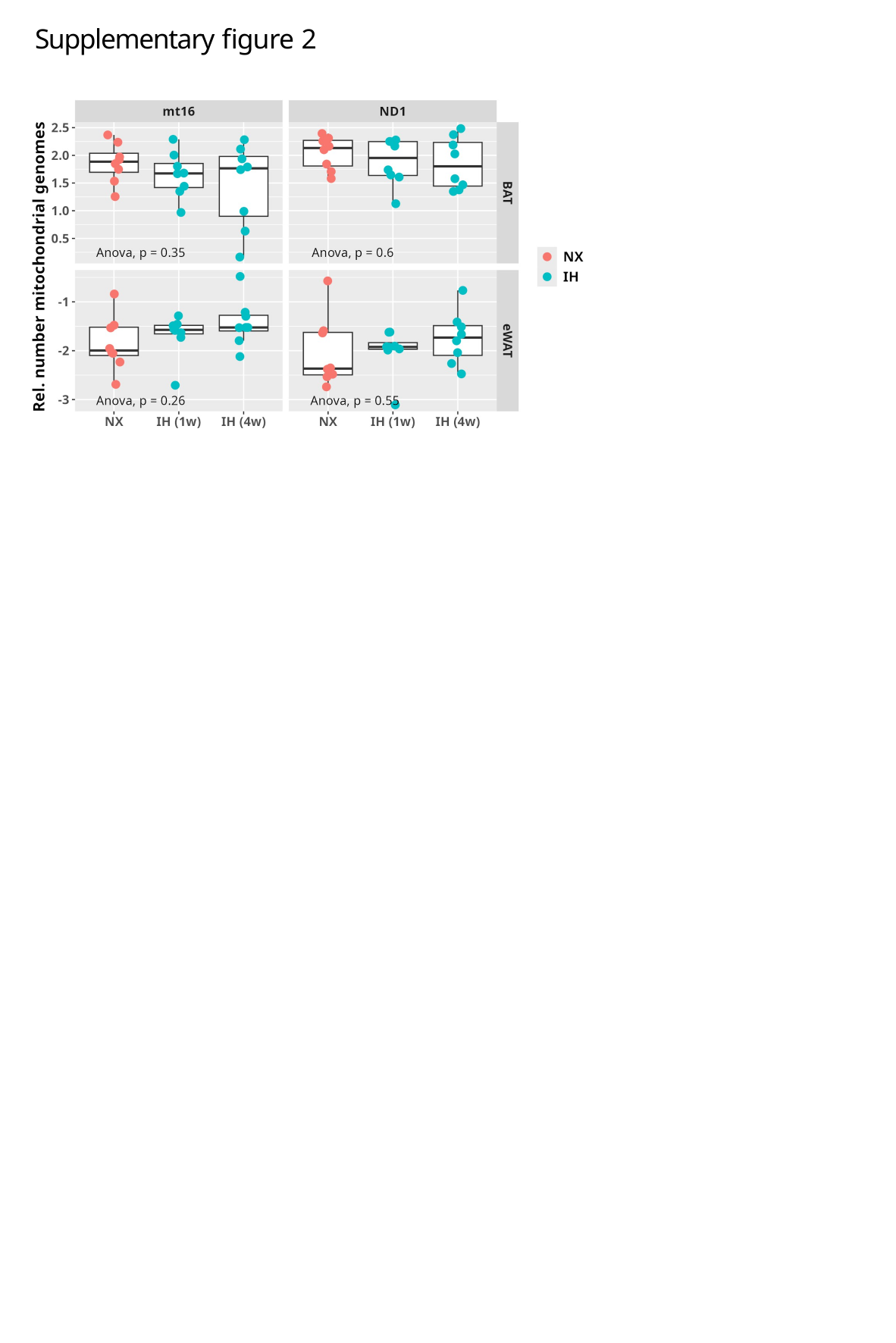

Supplementary figure 2

### Slide 3
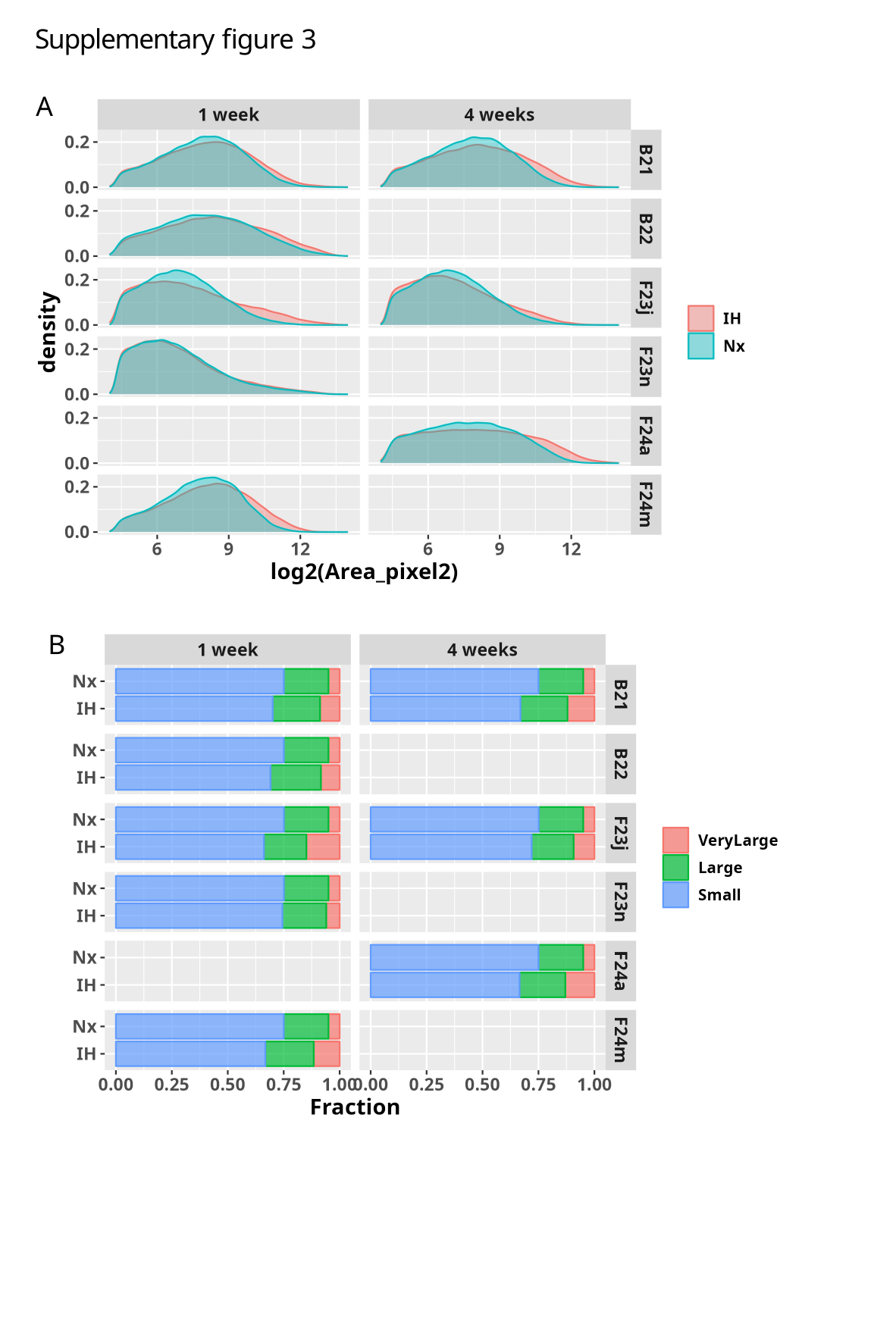

Supplementary figure 3
A
B

### Slide 4
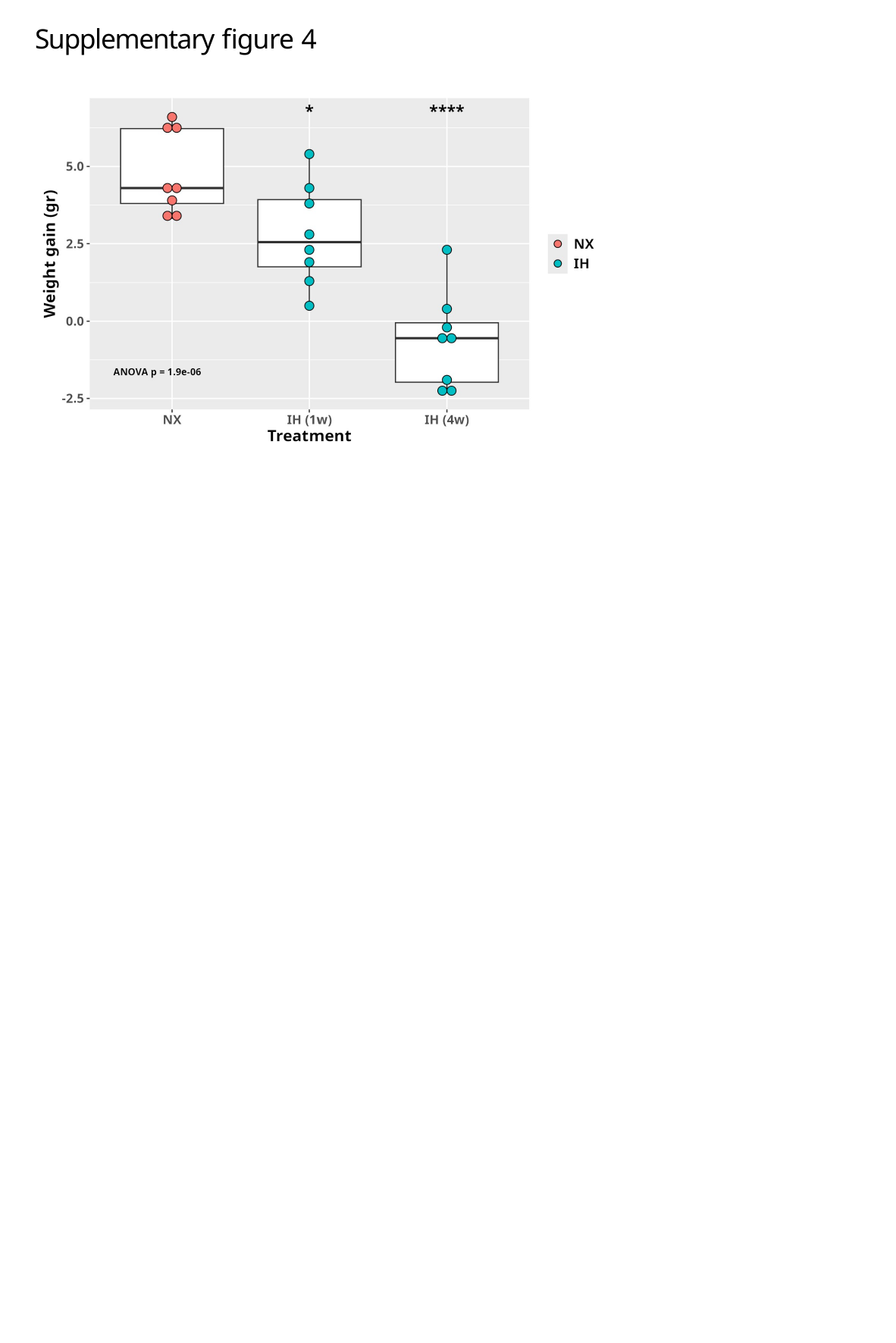

Supplementary figure 4
