## Supplementary material for "Intermittent Hypoxia Drives Early Metabolic Dysfunction in Brown Adipose Tissue": Text for supplemental Figures

### Supplementary figure 1. Correlation of transcriptomic responses over time.

Scatter plots comparing log₂ fold changes in gene expression (IH vs. normoxia) at 1 and 4 weeks in BAT, WAT, and liver. Only genes differentially expressed at both time points (FDR < 0.05; |log₂FC| > 0.5) are shown. The red line indicates linear regression; the blue line represents perfect correlation (slope = 1). Regression slope and R² values are provided.

### Supplementary figure 2. Mitochondrial DNA content.

Mitochondrial abundance was quantified by qPCR using primers for mt16 or ND1, normalized to nuclear gene Hk2. No significant differences were observed between groups (ANOVA, p > 0.05). Data are expressed relative to the mean of all WAT and BAT samples, illustrating lower mitochondrial DNA levels in WAT. On average, BAT had ~20- fold higher mitochondrial DNA than WAT.

### Supplementary figure 3. Effect of intermittent hypoxia on lipid droplet size in BAT.

(A) Lipid droplet size distributions in BAT across six independent experiments at 1 and 4 weeks of IH exposure. LDs were quantified as described in figure 3.
(B) LDs were classified into three size categories: small (<75th percentile), large (75th– 95th percentile), and very large (>95th percentile). Bar graphs show the proportion of each category across experiments and time points.

**Supplementary figure 4. Effect of intermittent hypoxia on boy weight gain.** Animals were treated as described in the Figure 2 legend. The graph shows the change in body weight for each animal, calculated as final weight minus initial weight. Statistical

analysis was performed using one-way ANOVA (F(2,21) = 26.31, p = 1.9 × 10⁻⁶), followed by post-hoc t-tests (*p < 0.05; **p < 0.0001).
